# The transcription factor FoxO1 is required for the establishment of the human definitive endoderm

**DOI:** 10.1101/2020.12.22.423976

**Authors:** Joshua Nord, Daniel Schill, Kirthi Pulakanti, Sridhar Rao, Lisa Ann Cirillo

## Abstract

The transcription factor FoxO1 has been shown to dynamically regulate cell fate across diverse cell types. Here, we employ a human induced pluripotent stem cell (hiPSC)-to-hepatocyte differentiation system that recapitulates the process of hepatocyte specification and differentiation in the human embryo to investigate FoxO1 as a participant in the molecular events required to execute the initial stages of liver development. We demonstrate that FoxO1 is expressed in hiPSC and at all stages of hepatocyte differentiation: definitive endoderm, specified hepatocytes, immature hepatoblasts, and mature hepatocyte-like cells. Disruption of FoxO1 activity by addition of the small molecule inhibitor AS1842856 at the beginning of the differentiation protocol abolishes the formation of definitive endoderm, as indicated by the loss of endoderm gene expression and the gain in expression of multiple mesoderm genes. Moreover, we show that FoxO1 binds to the promoters of two genes with important roles in endoderm differentiation whose expression is significantly downregulated in AS1842856 treated versus untreated cells. These findings reveal a new role for FoxO1 as an essential transcriptional regulator for the establishment of definitive endoderm in humans.

## Introduction

The transcriptional cues that give rise to definitive endoderm (DE) and subsequently derived tissues thereof have been widely studied, particularly in frogs, fish, and mice. Although several key transcription factors for DE differentiation have been identified, additional transcription factors are present and presumably active during this process with as of yet unspecified roles. One such transcription factor is FoxO1, which has been previously shown to play essential roles in cell fate decisions in multiple tissues. During the establishment of the definitive endoderm, it is imperative that inhibitory PI3K signaling is suppressed [1, 2]; this renders FoxO1 unphosphorylated, nuclear, and therefore transcriptionally active [3], raising the question of FoxO1’s involvement in the molecular events leading to DE formation. This manuscript uncovers a heretofore unknown and essential role played by FoxO1 in the establishment of the human definitive endoderm.

DE specification is an initial step in gastrointestinal development, with cells of the DE lineage giving rise to the gut tube and associated organs, including the liver, pancreas, and gallbladder. Originally described as the innermost germ layer, the DE is arguably the lesser studied of the three primary germ layers, the DE, mesoderm, and ectoderm, that arise during gastrulation. The DE and mesoderm both arise from bipotential progenitor ‘mesendoderm’ cells located in the anterior portion of the primitive streak and emerge during gastrulation by migrating outwards from the anterior streak [4–6]. Several cues are known to inform mesoderm vs endoderm cell fate. Of particular importance is Nodal, a determining factor in defining the cell fate of mesendoderm cells. Lower levels of Nodal signaling drives mesoderm formation, while higher levels drive endoderm formation [7–9]. The anterior portion of the primitive streak is the source of Nodal, which correlates with the fact that the DE is established in very close proximity. Nodal expression is activated and maintained by canonical Wnt signaling [10]. During this time PI3K/AKT signaling, which is inhibitory to both the Nodal and Wnt signaling pathways, must be inhibited to promote DE formation [1, 2].

Nodal and Wnt signaling act cooperatively to promote expression of a network of DE transcription factors, primarily by acting through the SMAD and β-catenin proteins, respectively. These transcription factors include members of the GATA, SOX and forkhead domain (FOX) families, all of which play essential roles in DE establishment and serve as markers thereof [11]. Roles for GATA-4 and −6 in DE formation are species dependent and semi-redundant [12–15]. In humans GATA-4 and −6 are required for induction of DE [16] and the subsequent differentiation of endoderm derived tissues including liver, pancreas, lung, and intestine (reviewed in [17]). While additional studies have demonstrated the necessity of GATA-4 and −6 for DE specification in frogs and zebrafish, in mice the role of GATA factors is limited to regulating extraembryonic endoderm lineages ([18] and reviewed in [19]). In contrast to GATA-4 and −6, SOX17 and FoxA2 have highly conserved functions across multiple species and are generally viewed as markers of DE. Although the exact targets, both direct and indirect, as well as how SOX17 regulates gene transcription are still debated, SOX17 has been shown to be indispensable for the formation of DE [20–23]. Via interactions with β-catenin, SOX17 has been shown to activate FoxA2 [24, 25], a forkhead transcription factor that works in concert with GATA factors to drive transcription of endodermal genes and, during later hepatogenesis, liver genes through chromatin opening and remodeling [26, 27].

An additional forkhead factor recently reported to be present in DE is FOXO1 [28]. FoxO1 function is inhibited by PI3K/AKT-mediated phosphorylation, which triggers cytoplasmic retention of FoxO1 and the consequent inhibition of its transcriptional activity [29–33]. FoxO1 is nuclear and transcriptionally active in DE [28] where PI3K/AKT signaling is silenced. Like the GATA factors and FoxA2, FoxO1 functions as an initial chromatin binding and remodeling “pioneer” factor [34]. This unique characteristic places FoxO1 at the forefront when studying embryonic development, which requires the specific engagement and activation of genes informing germline and tissue specification within silent, compacted chromatin. As a transcriptional activator, FoxO1 regulates a vast number of genes in multiple tissues with important roles in protein homeostasis, metabolism [33, 35–38], proliferation [39, 40], apoptosis [41–44], resistance to oxidative stress [45, 46] and inflammation [47, 48]. Not surprisingly, FoxO1 has been shown to dynamically regulate cell fate across diverse cell types. The differentiation of hepatic stellate cells [39], cardiomyocytes [40], adipocytes [49–51], osteoblasts [52], and, most recently, pancreatic β-cells [28], as well as the maintenance of stem cell pluripotency [53], have been shown to require FoxO1 activity. These attributes of FoxO1, together with its expression in DE, raise the question of whether FoxO1 might act similarly to promote specification of the DE.

In order to investigate possible roles for FoxO1 in the establishment of the DE, we employed a hiPSC-driven differentiation system capable of recapitulating DE induction. We show that FOXO1 is expressed throughout the entire process of DE differentiation from hiPSC and that small molecule inhibition of FOXO1 activity during DE induction abolishes the establishment of the DE and the expression of endoderm markers. We also demonstrate binding of FOXO1 to promotors of genes with important roles in DE formation. Taken together, these findings support a role for FoxO1 as an essential transcriptional activator for the establishment of DE in humans.

## Materials and Methods

### Cell Culture

The SV7 human iPS cell line was received from the laboratory of Dr Stephen Duncan (Medical University of South Carolina). The cells were maintained and subjected to differentiation using similar protocols to those described [54]. Briefly, hiPSC colonies were maintained on StemAdhere (Cat # S2071-500UG, Primorigen, Madison, WI) coated plates and fed daily with mTeSR1, prepared as previously described [55]. and supplemented with 20% MEF-conditioned medium at 37°C, 4% O_2_/5%CO_2_ and passaged as needed. To prepare for differentiation, colonies were dissociated with accutase (Cat #: SF006, Millipore Sigma, St. Louis, MO), seeded onto Matrigel (Cat #: 354230, Corning, Corning, NY) coated plates and allowed to recover for up to 24 hours. Differentiation was initiated and carried out with daily feedings of the following medium and growth factors: Day 1-2-RPMI/B27minus insulin (Cat #: A18956-01 Invitrogen, Carlsbad, CA), 100ng/mL Activin A (Cat #: 338-AC, R&D Systems, Minneapolis, MN), 20ng/mL FGF2 (Cat #: PHG0023, Invitrogen) and 10ng/mL BMP-4 (Cat #: 314-BP, Peprotech, Rocky Hill, NJ); Day3-5 – RPMI/ B27minus insulin, 100ng/mL Activin A; Day6-10 – RPMI/B27-with insulin-(Cat #: 17504-044, Invitrogen), 20ng/mL BMP-4 and 10ng/mL FGF2; Day11-15 – RPMI/B27-with insulin-, 20ng/mL Hepatocyte Growth Factor – HGF (Cat #: 100-39, Peprotech); Day16-20-Hepatocyte Culture Medium -HCM (Cat #: CC-3198, Lonza, Walkersville, MD) omitting the EGF and the addition of 20ng/mL Oncostatin-M (Cat #: 295-OM-050 R&D Systems).

### ASl842856-mediated Inhibition of FoxO1

Differentiations were carried out as described above. At 1-day post induction, fresh AS1842856 (Millipore Sigma) was added to the medium at a concentration of 0.1μM. DMSO was used at an equal volume in control differentiations.

### RNA Isolation and Quantitative RT-PCR

Total RNA was isolated from hiPSCs at given time points of hepatocyte differentiation using the QIAshredder and RNeasy mini kits (Qiagen, Germantown, MD). Complementary DNA (cDNA) was synthesized using MMLV Reverse Transcriptase (Cat #: 28025013, Invitrogen) along with dNTPs and random primers (Invitrogen). Using these cDNAs as templates, quantitative RT-PCR was completed using a CFX96 Real Time System (Bio-Rad, Hercules, CA) with specific primers at an annealing temperature of 64°C. The primers used for specific amplifications are as follows: OCT4: F: 5’-TCTCCCATGCATTCAAACTGAG-3’ R: 5’-CCTTTGTGTTCCCAATTCCTTC-3’ SOX2: F: 5’-CCCACCTACAGCATGTCCTACTC-3’ R: 5’-TGGAGTGGGAGGAAGTAAC-3’ SOX17: F: 5’-AGAATCCAGACCTGCACAAC-3’ R: 5’-GCCGGTACTTGTAGTTGGG-3’ FOXA2: F: 5’-TCAACGACTGTTTCCTGAAGGTGC-3’ R: 5’-TTCTCGAACATGTTGCCCGAGTCA-3’ HHEX: F: 5’-CATTTAGCGCGTCGATTCTG-3’ R: 5’-GATTCTCCAACGACCAGACC-3’ FGF17: F: 5’-CAACTCTACAGCAGGACCAG-3’ R: 5’-CTCACTCTCAGCCCCTTTG-3’ FOXO1: F: 5’-GCAGATCTACGAGTGGATGGTC-3’ R: 5’-AAACTGTGATCCAGGGCTGTC-3’ T: F: 5’-CCGACTCGCCCAACTTC-3’ R: 5’-CCCAACTCTCACTATGTGGATTC-3’ GATA4: F: 5’-AGATGGGACGGGTCACTATC-3’ R: 5’-CAGTTGGCACAGGAGAGG-3’. Each gene was assayed in 3 to 5 independent differentiations and ß-Actin was used for normalization. Expression units were determined using 2^-ΔCT^. Error bars represent SEM and p values were determined by 2-sample students T test (unpaired, two tailed).

### Immunostaining

Cells were fixed using fresh 4% paraformaldehyde in PBS. Cells were permeabilized with 0.4% Triton X-100 prior to blocking in a solution containing 3% BSA in PBS. Primary antibodies were applied in a 1% BSA solution in PBS overnight at 4°C. Secondary antibodies were subsequently applied for 1 hour at room temperature and DAPI was used as a counterstain. Micrographs were captured using an Eclipse TE300 fluorescent microscope (Nikon) and SpotCamera software. Images collected from experimentally treated samples and controls were identically processed. Primary antibodies directed against the following were used: OCT3/4 [sc-9081, lot L2211 (Santa Cruz Biotechnology, Dallas TX)], Sox17 [AF1924 lot (R&D Systems)], FoxA2 [H00003170-M12, lot DC121-6C12 (Novus Bio, Littleton, CO)], HNF4α [sc6556, lot I1211 (Santa Cruz Biotechnology)], AFP [A8452 (Sigma)], and Albumin [CL2513A (Cedarlane)]. The following secondary antibodies were used: Alexa-fluor anti-goat 568nm [A11057, lot 1010042 (Molecular Probes, Eugene, OR)), Alexa-fluor anti-rabbit 488nm [A21206, lot 1754421 (Invitrogen)], and Alexa-fluor anti-mouse 488nm [A21202, lot 1796361 (Invitrogen)].

### Western Blotting

Immunoblotting was performed as previously described [56]. Briefly, cells were lysed in RIPA buffer containing 150mM NaCl, 50mM Tris, 1% IGEPAL, 0.1% SDS and protease inhibitor cocktail (Millipore Sigma). These cell extracts were run on a 10% acrylamide gel before being transferred to a PVDF membrane (Bio-Rad). The membranes were blocked in 5% milk prior to incubation with primary antibodies overnight at 4°C. Washes were then conducted with TBST buffer prior to incubation with HRP conjugated secondary antibodies for two hours. The membrane was washed again prior to exposure to ECL Western blot detection reagent (GE Healthcare, Chicago, IL) and GeneMate Blue Autoradiography Film (VWR, Radnor, PA). Primary antibodies used were anti-FoxO1 [C29H4, lot 11 (Cell Signaling, Danvers, MA)] at a 1:1,000 dilution and anti-tubulin [sc-E-19-R, lot C1516 (Santa Cruz Biotechnology)] at a 1:1,000 dilution. Secondary antibody was goat anti-rabbit IgG-HRP at a 1:5,000 dilution [sc-2030, lot F2613 (Santa Cruz Biotechnology)].

### RNA-Seq Analysis

SV7 hiPSCs were subjected to hepatocyte differentiation out to day 3. Cells were treated for 48 hours with either AS1842856 or DMSO, as control, starting at day 1 post induction. Total RNA was extracted using TRIzol (ThermoFisher) and further processed via the RNeasy mini kit (Qiagen). RNA was collected from 3 independent differentiations. RNA quality was validated using an RNA ScreenTape on a TapeStation (Agilent Technologies, Santa Clara, CA). RNA concentration was determined by Nanodrop and 1μg of total RNA was used for library preparation. mRNA isolation was completed using NEBNEXT Poly(A) mRNA Magnetic Isolation Module (New England Biolabs, Cambridge, MA). Libraries were prepared using NEBNext Ultra RNA Library Prep Kit for Illumina (New England Biolabs) and NEBNext Multiplex Oligos for Illumina (New England Biolabs) according to the manufacturer’s specifications. Average library length was determined using a DNA ScreenTape on a TapeStation. Library concentration was determined by NEBNext Library Quant Kit for Illumina (New England Biolabs). Paired end sequencing (76 cycles) on libraries was performed on a NextSeq 500 (Illumina, San Diego, CA).

Raw sequences (FASTQ files) were aligned to hg19 using STAR [57]. Differential expression was performed using DE-seq [58]. Gene level absolute expression was quantified using CuffLinks [59]. Default parameters were used for all analyses unless otherwise specified.

### Chromatin Immunoprecipitation

SV7 hiPSCs were subjected to the first 3 days of the hepatocyte differentiation protocol, crosslinked with 1% formaldehyde for 10 minutes and quenched with 0.175M glycine. The cells were lysed in ChIP cell lysis buffer containing 0.5% IGEPAL, 85 mM KCl, 5 mM Pipes and 15 mM sodium butyrate, followed by nuclear lysis in 50 mM Tris-HCl pH 8.0, 10 mM EDTA, 1% SDS and 15 mM sodium butyrate. The chromatin was sonicated using a Bioruptor Pico (Diagenode, Denville, NJ) sonication device for 30 seconds on/30 seconds off for a total of 3 cycles or a Branson cell disruptor 185 sonifier at setting 4 for four pulses and 20 s/pulse with 2 minutes between pulses, which gave an average DNA fragment size of approximately 500 bp. The chromatin concentration was determined using a Bradford protein assay (Bio-Rad). The chromatin was diluted 1:6 in IP dilution buffer (1.1 % Triton X-100, 0.01 % SDS, 1.2 mM EDTA, 16.7 mM Tris-HCl pH 8.0, 16.7 mM NaCl, and 15 mM Na butyrate) to reduce the SDS concentration to amenable range for IP. Up to 300μg of chromatin was used for immunoprecipitation or set aside for total input. Antibodies directed against FoxO1 [ab39670, lot GR72068-47 (Abcam, Cambridge, MA)] or IgG [12-371B, lot 2713276 (Millipore Sigma)] were added to the chromatin and rotated overnight at 4°C. 50μL of pre washed Magna ChIP protein A+G magnetic beads (Millipore Sigma) were then added to the samples and rotated at 4°C for 2 hours to elute the chromatin:antibody complexes. Magnetic beads were isolated via a DynaMag-2 magnetic rack (Invitrogen). Sequential washes were completed with low salt buffer (2 mM EDTA, 20 mM Tris pH 8.0, 150 mM NaCl, 1 % Triton X-100, 0.1 % SDS), high salt buffer (2 mM EDTA, 20 mM Tris pH 8.0, 500 mM NaCl, 1 % Triton X-100, 0.1 % SDS), lithium chloride buffer (1 mM EDTA, 10 mM Tris pH 8.0, 1 % IGEPAL, 1 % w/v Na deoxycholate, 250 mM LiCl) and TE buffer (1 mM EDTA, 10 mM Tris pH 8.0). The DNA was eluted from the beads using freshly prepared elution buffer (100mM NaHCO3, 1% SDS) and the samples were incubated overnight at 67°C to reverse crosslinks. Purification of the DNA was performed using the Qiaquick PCR purification kit (Qiagen). Analysis was completed by quantitative PCR analysis of regions in the promoters of the selected genes. Primer sequences are as follows: APELA-5’: F: 5’-ACTGGGAGATGAACTAAGACTTG-3’ R: 5’-AAGCCACAAACTCATGAAATCTG-3’ LINC00261-5’: F: 5’-TTTCAGCCTCCATTGTCCC-3’ R: 5’-GAACACAAGGAAAAATTGGG-3’ LINC00261-3’: F: 5’-CAGAGACTTTCCCGACCATTC-3’ R: 5’-CCTCCACCCTCAACTTCATTG-3’. Percent total input was calculated. Error bars represent SEM and p values were determined by 2-sample students T test (unpaired, two tailed).

### ML221-mediated inhibition of APLNR

Differentiations were carried out as described above. At the onset of differentiation, fresh ML221 (Cat #: SLM0919-5MG, Millipore Sigma) was added to the medium at a concentration of 20μM. DMSO was used at an equal volume in control differentiations.

## Results

### FoxO1 is expressed throughout DE specification and human hepatocyte differentiation from hiPSC

The role(s) of FoxO1 in human DE formation is completely unexplored. To investigate this, we used a hiPSC-to-hepatocyte differentiation system that faithfully recapitulates the sequential steps of hepatocyte differentiation, including the initial step of DE specification, that occurs during *in vivo* development of the human embryonic liver (Figure 1A). The latter stages are documented by our immunofluorescent (IF, Figure 1A) and qRT-PCR (Figure 1B) analyses of marker protein and mRNA expression respectively, confirming the synchronous and developmentally accurate differentiation of hepatocytes from hiPSCs in our hands; importantly, 70-80% of the cells consistently express differentiation markers specific for each stage of human embryonic/fetal liver development, including DE formation. Importantly, as shown in Figure 1C and D, FoxO1 mRNA and protein, respectively, are expressed in the DE and all subsequent stages of hepatocyte differentiation. These data demonstrate the utility of the hiPSC driven hepatocyte differentiation system to examine the potential involvement of FoxO1 in DE formation.

**Figure 1.**
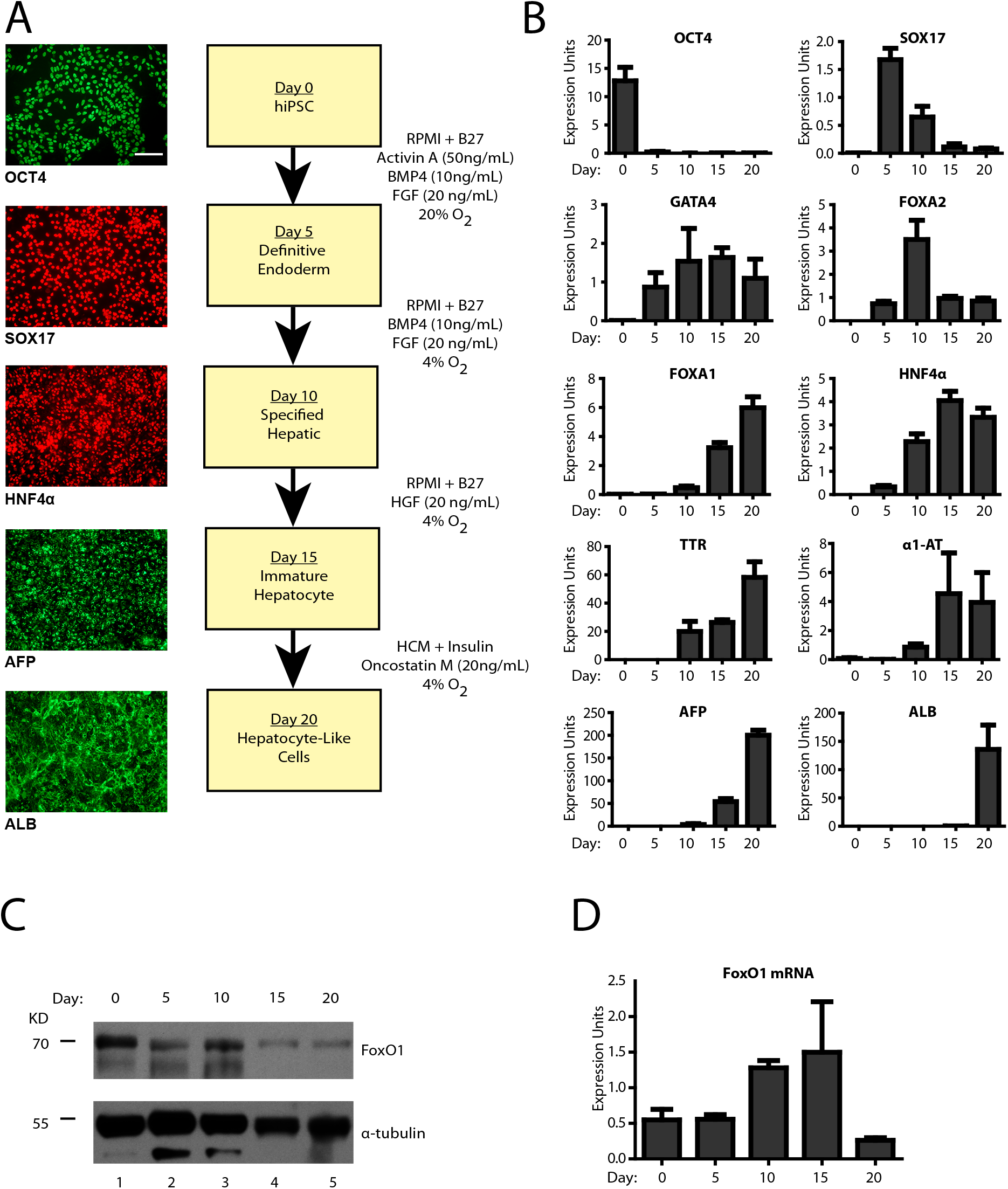
FoxO1 is expressed throughout hiPSC-directed hepatocyte differentiation. (A) The hiPSC-to-hepatocyte differentiation system with timeframes for stages of differentiation. Medium conditions for each stage are listed to the right. Representative immunofluorescent (IF) staining for stage specific markers, Oct4, SOX17, HNF4α, alpha-fetoprotein (AFP) and albumin, are displayed on the left. Images are at 20x magnification and the scale bar represents 100μm. (B) Western blot and (C) RT-qPCR analysis of FoxO1 expression during hepatocyte differentiation from hiPSC. Error bars represent the standard error of the mean (SEM) from 3 biologically independent differentiations

### Small molecule inhibition of FoxO1 activity abolishes the establishment of DE from hiPSC

To address the possible contribution(s) of FoxO1 to DE formation, we used the FoxO1 small molecule inhibitor AS1842856 [28, 50, 60–62]. Use of AS1842856 was necessitated by the fact that a knockout approach isn’t feasible due to the requirement of FoxO1 in hiPSC pluripotency maintenance [53]. AS1842856 inhibits FoxO1-mediated transcription via its specific binding to FoxO1 (and minimally to other FoxO factors [60]), preventing FoxO1’s ability to bind chromatin and recruit the histone acetyltransferase p300 to the transcription complex [60, 61]. When added directly to the differentiation media at day 1 post induction (Figure 2A), AS1842856 inhibition of FoxO1 completely abolished induction of DE. This was demonstrated by the loss of transcripts corresponding to the DE markers SOX17, FGF17, FoxA2, and HHEX, downregulation of GATA-4 transcripts as assessed by qRT-PCR (Figure 2B), as well as the loss of SOX17 and FoxA2 proteins as assessed by IF staining (Figure 2C). The absence of mRNA (Figure 2C) or protein (Figure 2E) corresponding to the pluripotency markers Oct4 and Sox2 indicated that loss of DE marker expression in the AS1842856-treated cells were no longer pluripotent, while expression of the mesoderm marker Brachyury (Figure 2B) intriguingly indicated that they may have adopted a mesoderm fate. Treatment with AS1842856 did not significantly alter levels of FOXO1 at the transcript (Figure 2D) or protein level (Figure 2E). Taken together, this data suggests that FoxO1 is required for the establishment of DE from hiPSC and suggests a potential change in cell fate from endoderm toward mesoderm in response to AS1842856-mediated inhibition of FoxO1 activity during hiPSC directed DE differentiation.

**Figure 2.**
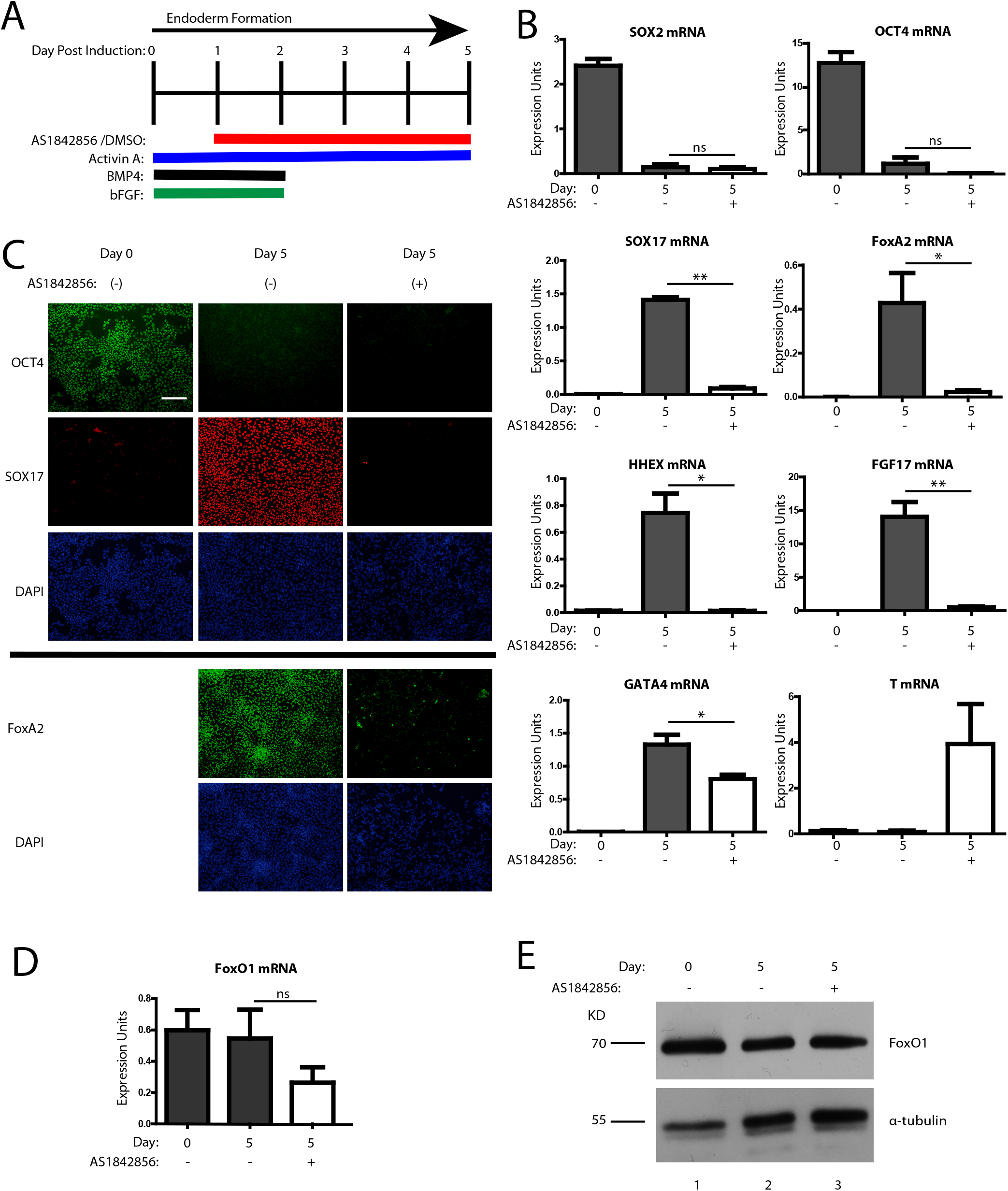
AS1842856-mediated inhibition of FoxO1 activity on day 1 of DE induction prevents DE specification. (A) Schematic depiction of hiPSC-directed DE differentiation and the AS1842856 treatment window. (B) RT-qPCR analysis of the indicated pluripotency, DE, and mesoderm markers in AS1842856 vs DMSO treated cells at day 5 post DE induction. Error bars represent the standard error of the mean (SEM) from a minimum of 4 biologically independent differentiations and significance was determined by student t-test, *<0.05 and **<0.01. (C) Representative IF analysis of day 0 (pluripotent) or AS1842856 vs. DMSO treated cells at day 5 post DE induction using antibodies directed against pluripotency marker Oct4 and the DE markers SOX17 and FoxA2. DAPI was used to identify nuclei. Images are at 20x magnification and the scale bar represents 100μm. (D) RT-qPCR analysis of FoxO1 transcript levels in AS1842856 vs DMSO treated cells at day 5 post DE induction. Error bars represent the standard error of the mean (SEM) from a minimum of 3 biologically independent differentiations. (E) Western blot analysis of FoxO1 protein in AS1842856 vs DMSO treated cells at day 0 (pluripotent) and day 5 (DE) post induction.

### AS1842856 treatment alters the transcriptional profile of cells generated from hiPSC-directed DE differentiation

FoxO1’s role as a transcriptional regulator led us to speculate that the misregulation of FoxO1 targets necessary for DE differentiation was responsible for the failure of DE formation observed in the presence of AS1842856. To investigate this possibility, we used RNA-seq to identify transcriptional targets of FoxO1 that are important for the differentiation of DE from hiPSC. In the hiPSC hepatocyte differentiation system used in this study, DE is fully established by day 5 post induction (Figure 1). We therefore chose to examine alterations in FoxO1-mediated gene expression at day 3 post induction, corresponding to the initial expression of stage specific markers for definitive endoderm (see SOX17, FGF17, HHEX in Table 1) and the likely time-point of global transcriptional changes resulting from FoxO1 loss. Comparison of gene expression profiles generated by RNA-Seq analysis of AS1842856 treated vs. DMSO treated cells harvested at day 3 post induction (Figure 3 A), revealed a total of 1904 genes whose expression was significantly altered (+/− ≥ 2-fold and corrected p< 0.05) by FoxO1 inhibition. Differential sequencing (DeSeq) analysis of the data showed that AS1842856 versus DMSO treatment resulted in two separate cell populations as illustrated by the heatmap depicting relative mRNA levels of the top 1000 differentially expressed genes shown in Figure 3B. Similarly, principle component analysis (PCA; Figure 3C) demonstrates the distinct grouping of transcripts corresponding to the AS1842856 versus DMSO treated differentiations.

**Figure 3.**
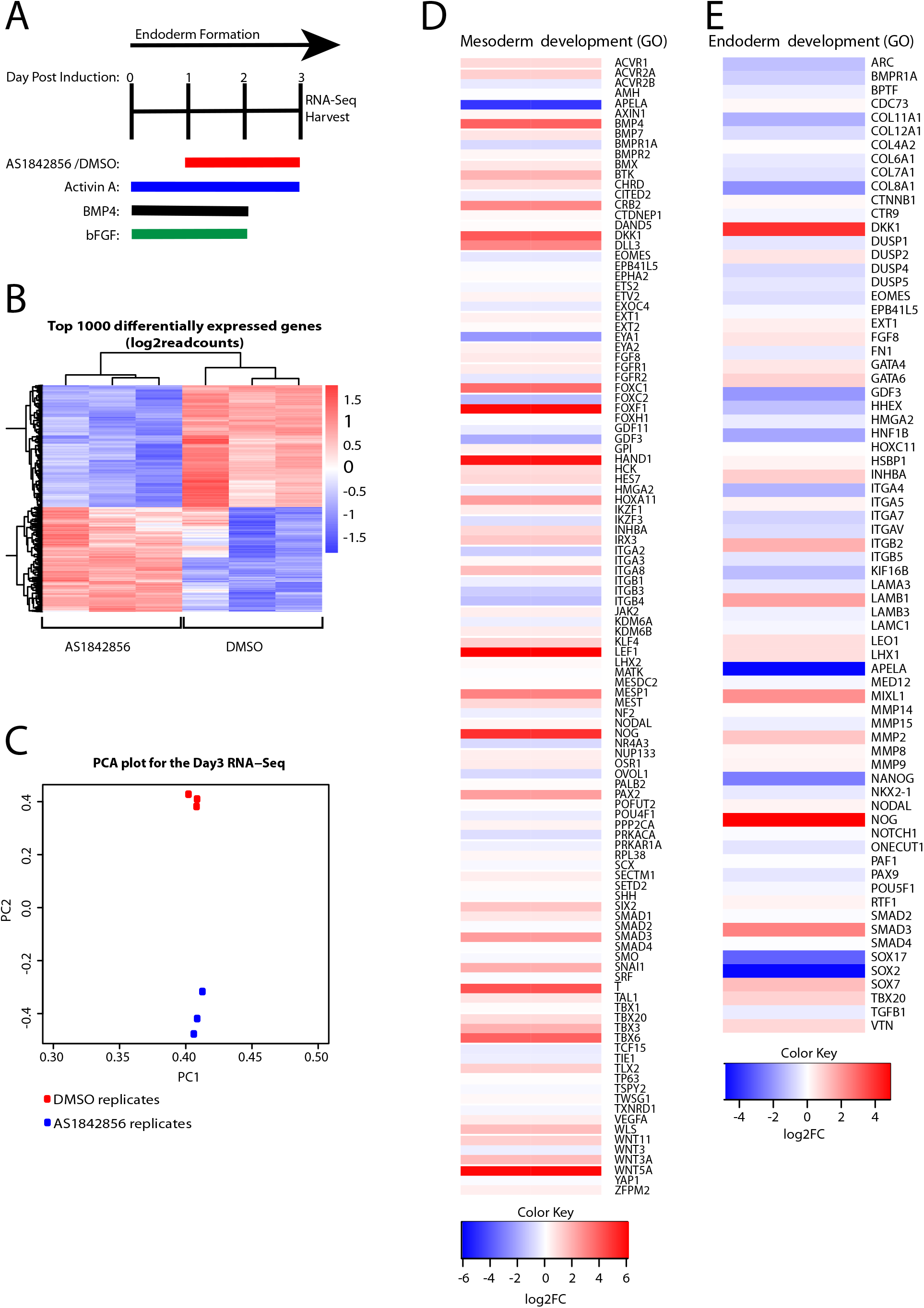
RNA-seq analysis reveals distinct transcriptional profiles in AS1842856 vs. DMSO treated cells harvested at day 3 post DE induction from hiPSC. Schematic depiction of hiPSC-directed DE differentiation, the AS1842856 treatment window, and the timing of RNA harvest. Cells harvested from 3 independent differentiations were used for the RNA-seq analysis, providing 3 biological replicates for AS1842856 and DMSO treatment. (B) Heatmap of gene expression profiles for the top 1000 differentially expressed genes in AS1842856 vs. DMSO treated cells at day 3 post DE induction. (C) Principle Component Analysis (PCA) of the variation in gene expression between AS1842856 vs. DMSO treated cell populations harvested at day 3 post DE induction. Each dot represents a biological replicate with red indicating the DMSO treated and blue indicating AS1842856 treated replicates. (D) Heatmap showing relative mRNA levels of mesoderm development genes in AS1842856 vs DMSO treated cells harvested at day 3 post DE induction. The mesoderm development gene set was retrieved from http://software.broadinstitute.org/gsea/msigdb/genesets.jsp. (E) Heatmap showing relative mRNA levels of the endoderm development genes in AS1842856 vs DMSO treated cells harvested at day 3 post DE induction. The endoderm development gene set was retrieved from http://software.broadinstitute.org/gsea/msigdb/genesets.jsp.

**Table 1.**
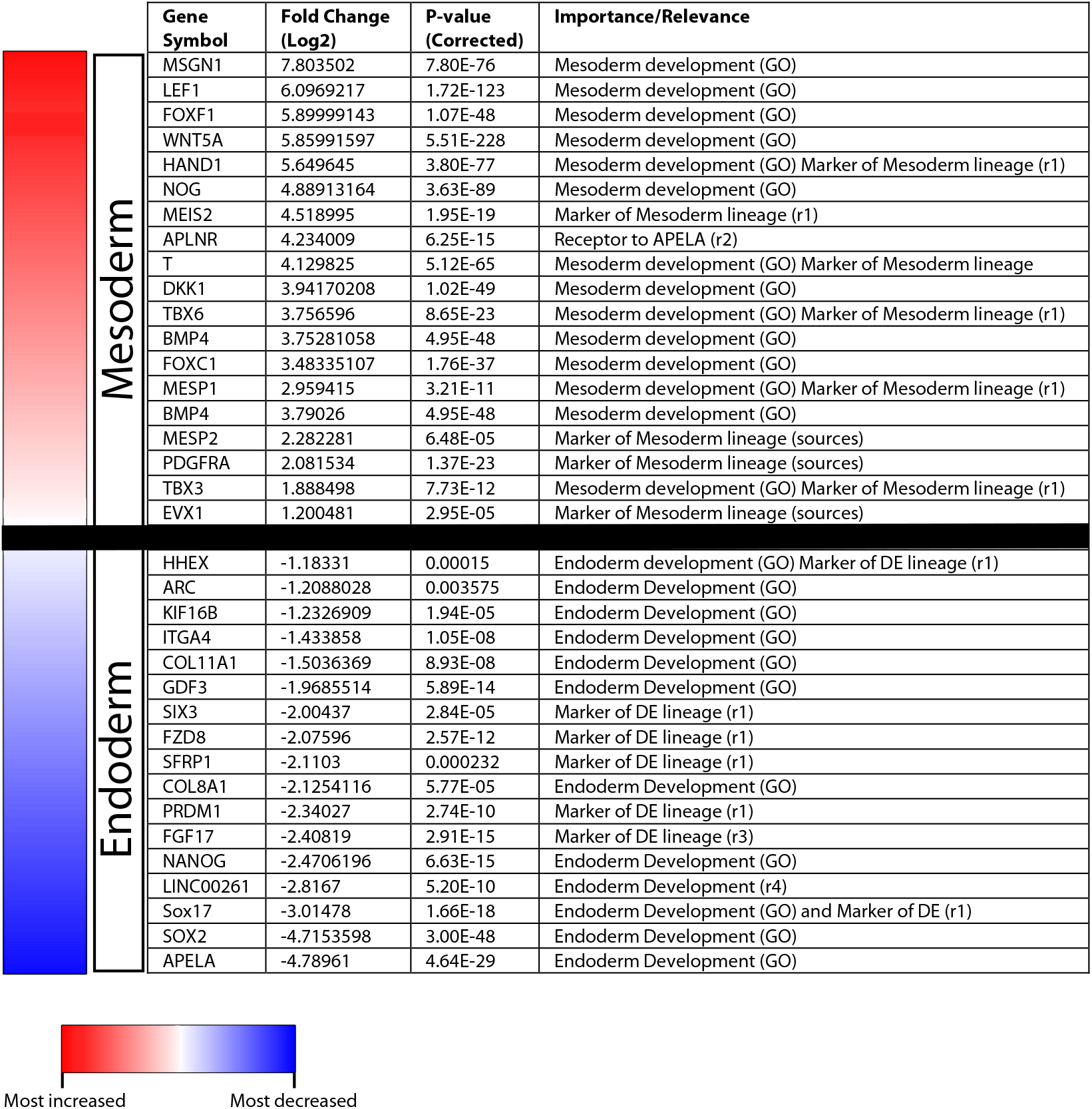
AS1842856 treated cells display an increase in mesoderm gene expression and a decrease in endoderm gene expression. This table depicts the expression levels of differentially expressed genes in AS1842856 vs. DMSO treated cells harvested at day 3 post DE induction with previously identified roles in mesoderm and endoderm development. Genes corresponding to transcripts exhibiting the biggest changes in expression in response to AS1842856 treatment with p-values under 0.01 were chosen from the Gene Ontology lists from the Broad Institute. Additionally, other genes were selected for this table, from the indicated sources, based upon their described roles in either the mesoderm or endoderm lineages. ‘GO’ identifies that the gene was obtained through GSEA analysis described in Figure 3 D-E. Other identifiers are as follows: ‘r1’:[63], ‘r2’:[69], ‘r3’:[65], ‘r4’:[64].

Utilizing the gene ontology (GO) gene sets provided by the BROAD institute, heatmaps were generated for genes pertaining to endoderm and mesoderm development. When comparing AS1842856 treatment to DMSO controls, it is apparent that many genes previously linked to mesoderm development are upregulated in response to FoxO1 inhibition (Figure 3D). The inverse is true for the endoderm development GO terms, where treatment with AS1842856 results in the reduction of many genes linked to endoderm development (Figure 3E). Table 1 shows a compilation of the top upregulated mesoderm genes and top downregulated endoderm genes with corrected p-values of less than .001 generated from the GO analysis shown in Figure 3 and from additional comparison of our RNA-seq data to that from recent publications [63–65]. A trend is readily apparent whereby cells generated from hiPSC in the presence of AS1842856 exhibit a transcriptional profile reflecting mesoderm in comparison to cells generated from hiPSC in the presence DMSO which exhibit a transcriptional profile reflecting DE. We conclude that AS1842856-mediated inhibition of FoxO1 abolishes DE formation while promoting aberrant differentiation toward a mesoderm fate.

### AS1842856 treatment during DE differentiation from hiPSC inhibits expression of FoxO1 target genes with important roles in DE differentiation

In order to identify FoxO1 target genes with roles in DE formation, we used JASPER [66] to search for FoxO1 consensus sites in the regulatory regions of the endoderm genes shown in Table 1 whose expression is significantly downregulated by AS1842856 treatment. This search revealed presumptive FoxO1 binding sites within the promoters of two genes, LINC00261 (DEANR1) and APELA, having proven roles in definitive endoderm differentiation [64, 67, 68]. As shown in Figure 4A, the DEANR1 promoter contains one FoxO1 consensus site at 275 bp upstream of the transcriptional start site (TSS), while the APELA promoter contains two FoxO1 consensus sites, at 310 bp and 598 bp upstream of the TSS. The long non-coding RNA DEANR1 is essential for SMAD recruitment to the FoxA2 promoter; not surprisingly, knockout of DEANR1 in human ES cells diminishes their ability to differentiate into DE [64]. APELA, a ligand of the G-coupled protein receptor APLNR [69], has been shown to play a role in endoderm formation and maintenance [67, 68, 70]; it has been speculated to act in this capacity through regulation of Sox17 and FoxA2 (reviewed in [71]). Knockout of APELA in zebrafish causes defects in endoderm formation [68], while studies in human embryonic stem cells (hESC) demonstrate that APELA is capable of priming hESC towards DE by activating the TGF-ß pathway [67]. APELA displayed the greatest decrease in transcript expression in response to AS1842856-mediated inhibition of FOXO1. Moreover, APELA’s receptor, APLNR, which was recently shown to exhibit DE-enriched expression [16], is dramatically upregulated in this context, a common response to downregulation of GPCR ligands (Table 1). It should be noted that while APELA appears on both the mesoderm development and endoderm development GO gene set lists, its role in mesoderm development is non-cell autonomous.

The presence of FoxO1 binding sites in the DEANR1 and APELA promoters suggested that FoxO1 might promote DE formation, in part, through direct transcriptional activation of the DEANR1 and APELA genes. To address this, we performed chromatin immunoprecipitation (ChIP) analysis of FoxO1 binding to the DEANR1 and APELA promoters in DE at 3 days post induction from hiPSC (Figure 4A). ChIP confirmed FoxO1 binding to both the DEANR1 and APELA promoters (Figure 4B). In contrast, FoxO1 binding was not observed to a 3’ region of the DEANR1 gene that does not contain a FoxO1 consensus site. IgG does not immunoprecipitate chromatin corresponding to any of the regulatory regions examined. We conclude that FoxO1 binds to the promoters of two genes with essential roles in DE formation whose expression is lost in response to inhibition of FoxO1 activity.

### Small molecule inhibition of the APELA receptor APLNR results in decreased expression of key markers of DE differentiation

While knockout of DEANR1 has been shown to abolish DE differentiation from human ES cells, the impact of APELA on hiPSC-directed DE differentiation is unknown. To assess the role of APELA in DE differentiation from hiPSC, we inhibited the activity of its receptor, APLNR. hiPSC were differentiated to the day 5 DE stage in the presence of the APLNR small molecule inhibitor ML221 [72], or DMSO as a control (Figure 5A). As shown in Figure 5B, ML221 treatment resulted in significant reductions in the transcript levels of the DE markers Sox17 and FGF17 (Figure 5B). Although not statistically significant, a downward trend in FoxA2 transcript levels was also observed, whereas HHEX transcription was unaffected. We conclude that inhibition of the APELA receptor APLNR curtails expression of essential genes for DE formation. This finding suggests that the negative impact of the inhibition of FoxO1 activity on DE formation could be mediated, in part, by the consequent reduction in levels of APELA.

**Figure 4.**
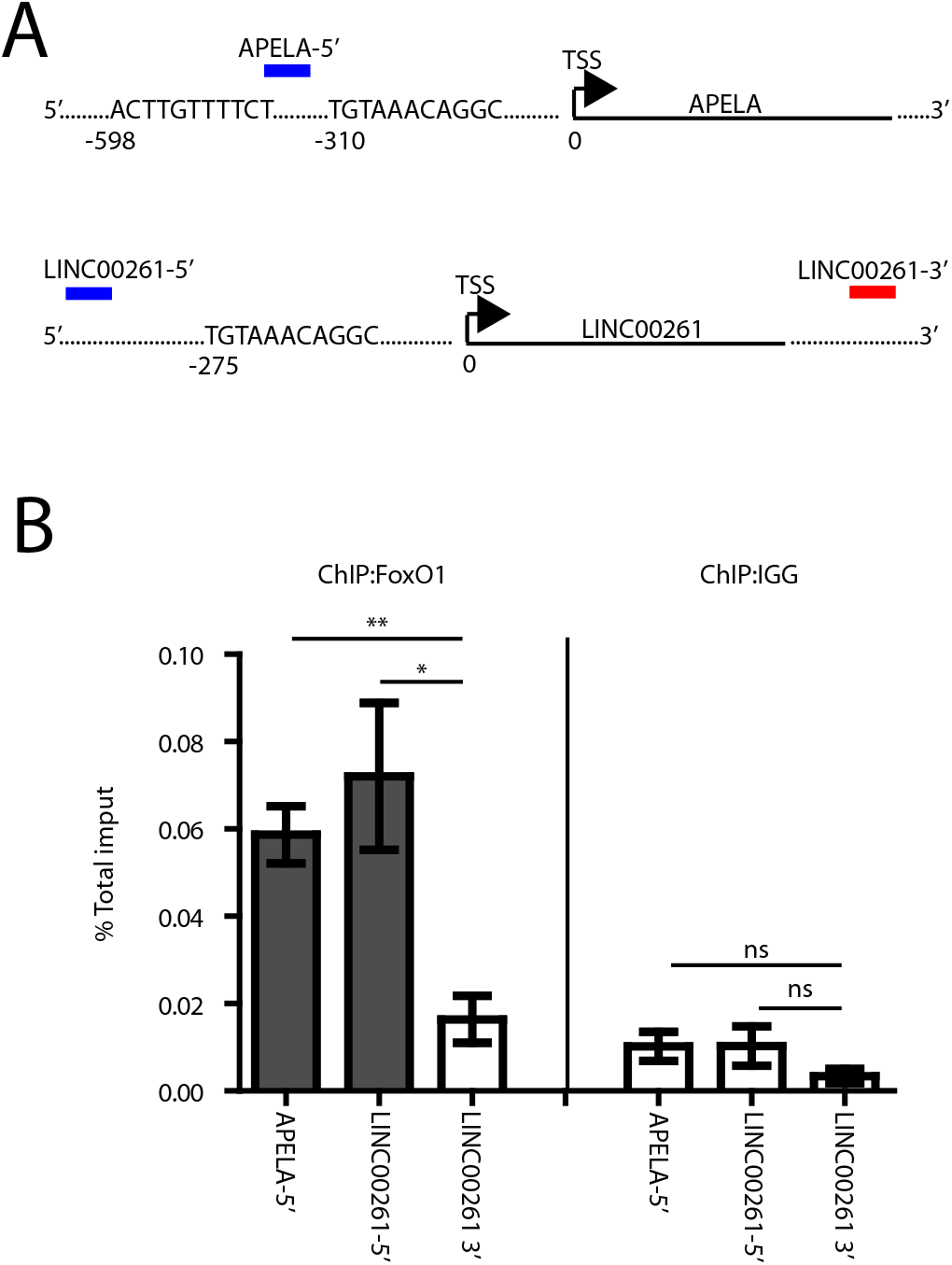
FoxO1 binds to the promoters of the endoderm genes APELA and DEANR1. (A) Schematic depiction of APELA and DEANR1 showing the predicted FoxO1 binding sites within their promoter regions. The transcriptional start sites (TSS) are labeled and marked as 0. Negative numbers are used to identify the distance of the presumptive FoxO1 binding sites from the TSS. The approximate location of the primers used for the ChIP-qPCR analysis of each promoter (blue bars) and the DEANR1 3’ region (negative control, red bars). (B) ChIP-qPCR analysis of FoxO1 binding to chromatin isolated from cells at day 3 post DE induction using antibodies against FoxO1 and, as a control, IgG. Values are expressed as % total input. Error bars represent the SEM from 4 independent differentiations and significance was determined by student t-test, *<0.05 and **<0.01.

**Figure 5.**
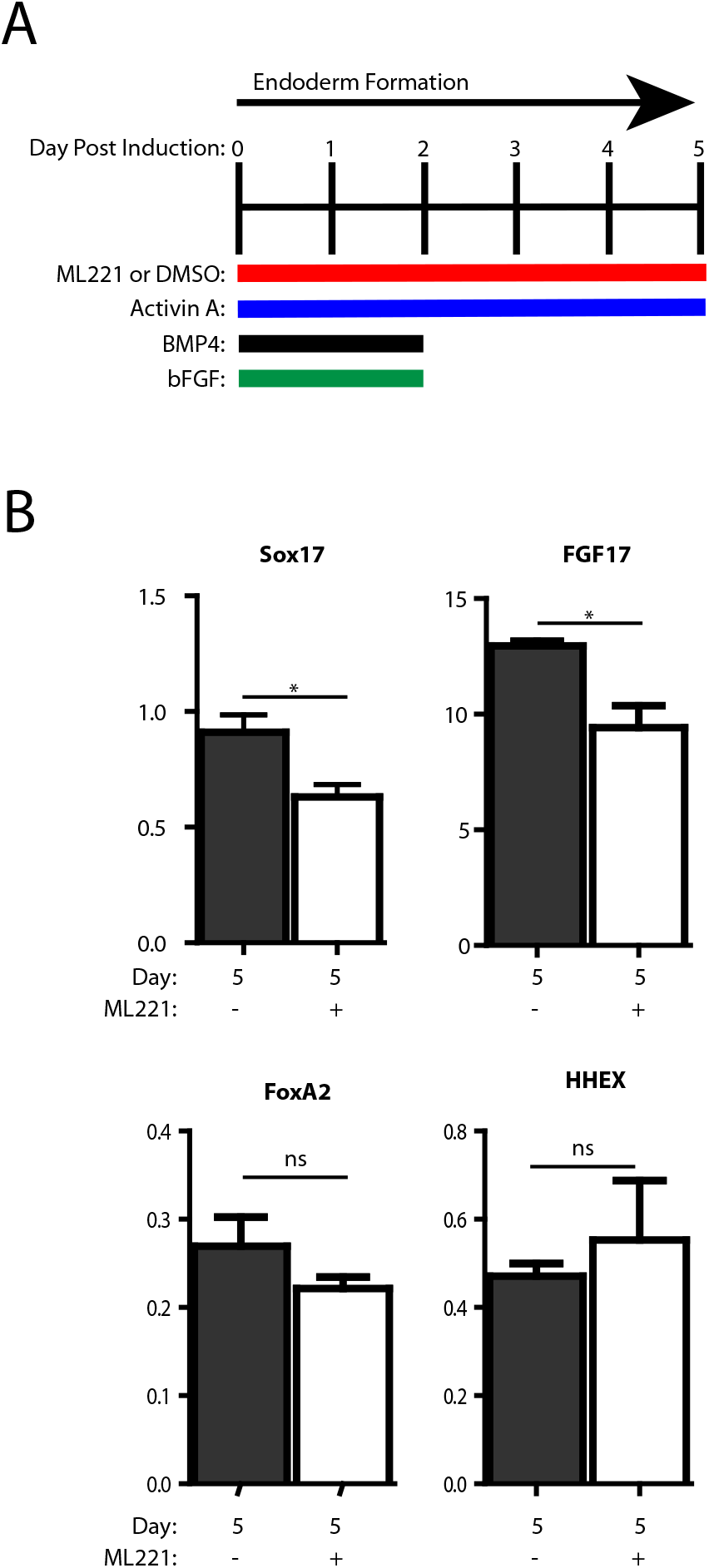
Treatment with the APLNR inhibitor ML221 results in reduced expression of several markers of the DE lineage. (A) Schematic depiction of hiPSC-directed DE differentiation and the ML221 treatment window. (B) RT-qPCR analysis of the endoderm markers SOX17, FGF17, FoxA2 and HHEX. Values are expressed as expression units. Error bars represent the SEM from 4 individual differentiations and significance was determined by student t-test, *<0.05.

## Discussion

This study demonstrates, for the first time, that FoxO1 plays a pivotal role as a transcriptional activator during human DE formation. Using a directed hiPSC-to-hepatocyte differentiation system we showed that the inhibition of FoxO1 activity using the small molecule inhibitor AS1842856 abolishes DE formation (Figure 2). The involvement of FoxO1 as a transcriptional activator in the context of differentiation and development is not new, as FoxO1 has been shown to transcriptionally regulate genes required for adipocyte, muscle, and osteoblast differentiation [40, 49–52]. FoxO1 has long been shown to be important in embryonic development, with FoxO1 null mice dying in utero [73–75]. FoxO1 null embryos display vascular and cardiovascular malformations, possibly linked to placental dysmorphogenesis and progenitor cell death (reviewed in [76]). Additionally, FoxO1 has been shown to be involved in the very earliest developmental processes through its ability to regulate hESC pluripotency through transcriptional regulation of the pluripotency associated genes Oct4 and Sox2 [53]. Contrary to the demonstrated role for FoxO1 in maintaining pluripotency of hESC, FoxO1 null mice show no pre-gastrulation defects [74, 77]. One explanation for this apparent contradiction is the redundancy of FoxO family members in mice that may not reflect the situation in humans. For example, unlike in human ES cells, pluripotency of mouse ES cells is controlled by both FoxO1 and FoxO3 [53, 78]. This redundancy of FoxO1 and FoxO3 in early mouse development, together with a prior study that showed FoxO1 to be virtually undetectable in fetal liver by northern blot until mouse embryonic day 18 [79], likely contributed to FoxO1’s remaining unstudied in DE or embryonic liver human liver development. It is therefore imperative to consider species-specific differences in FoxO redundancy when comparing roles for FoxO factors in development of mice versus humans.

RNA-seq analysis of AS1842856 vs DMSO treated cells at day 3 post DE induction demonstrated that AS1842856 mediated inhibition of FoxO1 activity causes increased expression of mesoderm markers and reduced expression of many endoderm markers, including Sox17, FGF-17, and HHEX (Figure 3 and Table 1). During gastrulation, a bipotential population of mesendoderm cells has the potential to adopt either a mesoderm or DE fate [4]. The fate switch from endoderm to mesoderm observed in response to AS1842856 inhibition of FoxO1 activity is therefore not surprising. Although more research is required, our data suggests a model whereby the mesendoderm population present during the initial stages of hiPSC-driven DE differentiation defaults to a mesoderm fate in the absence of FoxO1-mediated activation of genes required for DE formation. JASPER analysis for the FoxO1 consensus site identified two endoderm expressed genes, DEANR1 and APELA, containing one or more FoxO1 binding sites in their promoter. ChIP experiments performed on cells at day 3 post DE induction showed that FoxO1 binds to the promoter regions of both DEANR1 and APELA, providing strong evidence that FoxO1 acts as a transcriptional activator for both of these genes in DE. While the contribution of DEANR1 in DE formation through FoxA2 activation is well understood [64], that of APELA is less clear. While APELA knockout mice do not present an endoderm defect [80], APELA is required for differentiation of DE precursors in zebrafish, where it is postulated to enhance Nodal signaling [70] necessary for the specification of the DE. A similar scenario has been reported for DE lineage commitment from mesendoderm during human ES cell-directed DE differentiation [67]. APELA has been shown to function independently of the APLNR receptor in some contexts. For example, APELA’s role as a pluripotency factor in human ES cells does not require APLNR [67]. This suggests that APELA likely targets as of yet unknown receptors to carry out additional functions outside of its ability to regulate processes downstream of APLNR. Therefore, the modest, though statistically significant, reductions in the expression of the DE markers SOX17 and FGF17 observed in response to ML221 inhibition of APLNR may not reflect the entirety of the impact of APELA on DE formation.

In summary, we conclude that FoxO1 plays an important role as a transcriptional regulator for the establishment of DE, in part through its direct activation of the DEANR1 and APELA genes. Given the large number of additional genes whose expression is negatively impacted by inhibition of FoxO1 activity during hiPSC-driven DE formation, it is very likely that FoxO1 directly regulates multiple genes with roles in determining DE fate.

## Acknowledgements

We would like to thank Dr. Stephen Duncan (Medical College of South Carolina) for his kind gift of the SV7 hiPSCs and his lab’s generous instruction on hiPSC-directed hepatocyte differentiation. This work was supported by grants to LAC from the National Institutes of Health [DK093763, DK120548].

